# TENET: Gene network reconstruction using transfer entropy reveals key regulatory factors from single cell transcriptomic data

**DOI:** 10.1101/2019.12.20.884163

**Authors:** Junil Kim, Simon Toftholm Jakobsen, Kedar Nath Natarajan, Kyoung Jae Won

## Abstract

Accurate prediction of gene regulatory rules is important towards understanding of cellular processes. Existing computational algorithms devised for bulk transcriptomics typically require a large number of time points to infer gene regulatory networks (GRNs), are applicable for a small number of genes, and fail to detect potential causal relationships effectively. Here, we propose a novel approach ‘TENET’ to reconstruct GRNs from single cell RNA sequencing (scRNAseq) datasets. Employing transfer entropy (TE) to measure the amount of causal relationships between genes, TENET predicts large-scale gene regulatory cascades/relationships from scRNAseq data. TENET showed better performance than other GRN reconstructors, in identifying key regulators from public datasets. Specifically from scRNAseq, TENET identified key transcriptional factors in embryonic stem cells (ESCs) and during direct cardiomyocytes reprogramming, where other predictors failed. We further demonstrate that known target genes have significantly higher TE values, and TENET predicted higher TE genes were more influenced by the perturbation of their regulator. Using TENET, we identified and validated that Nme2 is a culture condition specific stem cell factor. These results indicate that TENET is uniquely capable of identifying key regulators from scRNAseq data.

**Key Points:**

- TENET measures putative causal relationships between genes using transfer entropy.
- TENET shows outstanding performance in identifying key regulators compared to existing methods.
- TENET can reveal previously uncharacterized regulators.

## INTRODUCTION

Regulatory mechanisms are key to understanding cellular processes. The cell-type specific functions and responses to external cues is governed by complex gene regulatory networks (GRNs) (Davidson and Levine 2008; Møller and Natarajan 2020; Kim et al. 2012). Various approaches including genome-wide location analysis using chromatin immunoprecipitation followed by genomewide sequencing (ChIP-seq) (Gerstein et al. 2012; Chen et al. 2008) and perturbation analysis were designed to explain the putative causal relationships between genes (Loh et al. 2006; Hormoz et al. 2016). However, protein binding information is limited by the availability of antibodies and identification of target genes is difficult when bound at intergenic regions. Moreover, using perturbation experiments, it is hard to measure the strength of the putative causal relationships with the target genes. Systems biology approaches have been suggested to predict regulators and their target genes, prior to experimental wet-lab validation to reduce the cost and time (Hartemink 2005; Zou and Conzen 2005; Margolin et al. 2006; Møller and Natarajan 2020; Cho et al. 2007). However, previous attempts to infer GRNs have been limited to a small number of genes (Moignard et al. 2015; Li et al. 2004; Sanchez-Castillo et al. 2018) and/or cannot detect putative causal relationships effectively (Aibar et al. 2017; Chan et al. 2017).

When dealing with causal relationships, time is often involved, i.e. an effect cannot occur before its cause. In order to utilize time to identify the cause (the regulator) and the effect (the target genes), a time series analysis of gene expression data would be useful. Single cell RNA sequencing (scRNAseq) provides sequential static snapshots of expression data from cells aligned along the virtual time also known as pseudo-time (Setty et al. 2016; Trapnell et al. 2014; Haghverdi et al. 2016). Indeed, gene expression patterns and peak levels across pseudo-time have been used to infer potential regulatory relationships between genes previously (Trapnell et al. 2014; van Dijk et al. 2018). It is based on an assumption that the expression profile of a potential regulator precedes the expression pattern of a target gene along the pseudo-time. Moreover, current approaches rely on visual and manual inspection and the gene expression dependencies are not extensively investigated. Systematic approaches that quantify potential causal relationships between genes and reconstruct GRNs are still highly required.

We aim to quantify the strength of causality between genes by using a concept originating from information theory, called transfer entropy (TE). TE measures the amount of directed information transfer between two variables (Schreiber 2000; Hlaváčková-Schindler et al. 2007). Leveraging the power to measure potential causality, TE has been successfully applied to estimating functional connectivity of neurons (Orlandi et al. 2014; Wollstadt et al. 2014; Spinney et al. 2017) and social influence in social networks (Kim et al. 2016). Based on TE, we developed TENET (https://github.com/neocaleb/TENET) to reconstruct GRNs from scRNAseq data. Using single-cell gene expression profile along the pseudo-time, TENET calculates TE values between each set of gene pairs.

We found that TE values of the known critical regulators (i.e. target genes) were significantly higher than that of randomly selected targets. Interestingly, target genes with higher TE values were influenced more profoundly by the perturbation analysis. We also show that TENET outperforms previous GRN constructors in identifying target genes.

Pseudo-time has been used in a number of GRN reconstructors (Matsumoto et al. 2017; Specht and Li 2017; Papili Gao et al. 2018; Qiu et al. 2020; Deshpande et al. 2019). Unique to TENET is the ability to represent key regulators with the hub nodes in the reconstructed GRNs. For instance, TENET identified pluripotency factors from scRNAseq during mouse embryonic stem cell (mESC) differentiation (Tuck et al. 2018) and the key programming factors from scRNAseq for the direct reprogramming toward cardiomyocytes (Liu et al. 2017b), where existing methods either failed to identify or capture their importance for the regulatory network. Interestingly, the factors that TENET identified were more negatively correlated with the number of final states (or attractors) in the Boolean networks (Moignard et al. 2015), which confirms the importance of the identified hub nodes. An alternative method SCENIC also infers GRNs and their target genes using co-expression and the motif information (Aibar et al. 2017). Compared with SCENIC, TENET determines the regulatory relationships using the expression profiles alone along the pseudo-time. Therefore, TENET can be used to search for any type of regulators regardless of their binding to DNA.

Applying TENET to scRNAseq data for mESC differentiation into neural progenitor cells (NPCs), we found that Nme2 is a key transcription factor (TF) that regulates pluripotency genes in a culture condition dependent manner. Inhibition of Nme2 in mESC cultured in ground state (2-inhibitors and LIF; ‘**2iL**’) conditions leads to dramatically reduced cell proliferation and reduced expression of its direct targets, compared to differentiation in permissive conditions (Serum and LIF; ‘**SL**’). In summary, TENET has a potential to elucidate previously uncharacterized regulatory mechanisms by reprocessing scRNAseq data.

## METHODS

### The TENET algorithms

TENET measures the amount of putative causal relationships using the scRNAseq data aligned along pseudo-time. From pseudo-time ordered scRNAseq data (Figure 1a), TENET calculates bidirectional pairwise TE values for selected genes using JAVA Information Dynamics Toolkit (JIDT) (Lizier 2014) (Figure 1b). We calculated TE values by estimating the joint probability density functions (PDFs) for mutual information (MI) using a non-linear non-parametric estimator “kernel estimator”(Schreiber 2000). The joint PDF of two genes *x* and *y* can be calculated as follows:

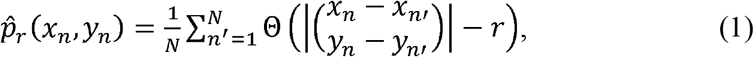

where Θ is a kernel function and *N* is the number of cells. We used step kernel (Θ(*x*>0)=1, Θ(*x*≤0)=0) with kernel width *r*=0.5 as default. The TE from *X* to *Y* is defined as follows:

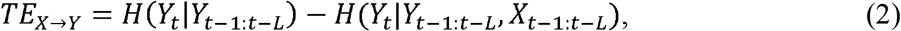

where *H*(*X*) is Shannon’s entropy of *X* and *L* denotes the length of the past events considered for calculating TE. It calculates the amount of uncertainty of *Y_t_* reduced by considering *X*_*t*-1:*t-L*_. We reconstructed the GRNs by integrating all TE values for gene pairs (Figure 1c). To remove potential indirect relationships, we applied the data processing inequality (Margolin et al. 2006), i.e. iteratively eliminating feed-forward loops. The feed-forward loop is defined by a network motif composed of three genes, where gene X regulates gene Y and both gene X and Y regulate gene Z. We trimmed the link from gene X to gene Z if TE_X_→_Z_ is less than the minimum value of TE_X_→_Y_ and TE_Y_→_Z_. Finally, we reconstructed a GRN consisting of the significant links using Benjamini-Hochberg’s false discovery rate (FDR) (Benjamini and Hochberg 1995) after performing the one-sided z-test while considering the all trimmed TE values as a normal distribution. The hub node is identified by calculating the number of targets (outgoing links).

**Figure 1.**
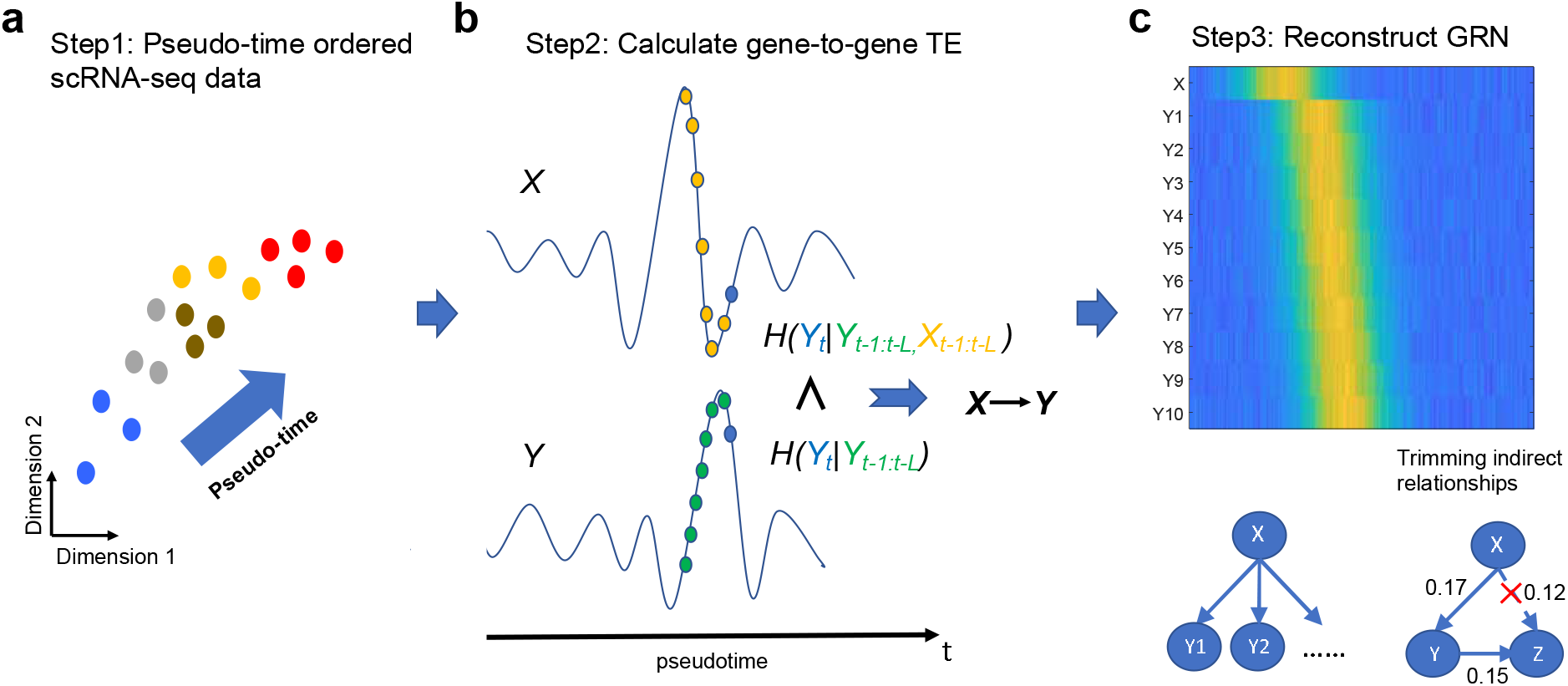
TENET reconstructs GRNs from pseudo-time ordered single cell transcriptome data using TE. **a**. Step 1: Pseudo-time ordered scRNAseq data are used as the input for TENET. **b**. Step 2: TENET calculates gene-to-gene pairwise TE while considering the past events of X and Y. **c**. Step 3: A reconstructed GRN is composed of putative but significant causal relationships followed by trimming indirect relationships. The heatmap shows the gene expression levels for a regulator (X) and its target genes.

### Statistical analysis

A two-sided one-sample z-test was performed to evaluate the mean of TE values for the targets of key factors (c-Myc, n-Myc, E2f1, Zfx, Nme2) in mESCs and Gata4 in mouse cardiomyocytes. This was accomplished by generating a fitted z-distribution of the TE values using the same number of randomly selected genes (1,000 times). A two-sided two-sample Student t-test was performed to evaluate the relative gene expression changes after knocking-in of Tbx3 and Esrrb and knocking-down of Pou5f1 and Nanog for the specified TE values, respectively.

### Data processing of scRNAseq data

To test TENET, we used the scRNAseq dataset obtained from mESCs (Tuck et al. 2018) and mouse cardiomyocytes (Liu et al. 2017b). Wishbone (Setty et al. 2016) was used for pseudo-time analysis. As an input gene list for the benchmarking of mESC dataset, we used 3,277 highly variable genes with log2(count)>1 for more than 10% of all cells and a coefficient of variation > 1.5. For the scRNAseq data during the reprogramming into cardiomyocytes, we used 8,640 highly variable genes with log2(count)>1 for more than 10% of all cells and a coefficient of variation >1. To reconstruct the GRN, we used a regulator gene list which includes genes with a GO term “regulation of transcription (GO:0006355)” for the mESC. We generated all the network figures using Cytoscape 3.6.1 (Shannon et al. 2003).

### Gene ontology (GO) terms and Kyoto Encyclopedia of Genes and Genomes (KEGG) pathways for the functional gene group

All enriched GO terms and KEGG pathways were obtained using Enrichr (Kuleshov et al. 2016). The “pluripotency gene” and the “neural differentiation gene” were obtained from the genes with a GO term “stem cell population maintenance (GO:0019827)” and “neuron differentiation (GO:0030182)”, respectively. We used GO terms “cardiac muscle cell differentiation (GO:0055007)”, “cardiac muscle contraction (GO:0060048)” for cardiomyocyte gene.

### Gene expression and ChIP-seq data for validation

We downloaded an RNAseq dataset in mESCs with three different combinations of double knock-in for Esrrb and Tbx3 (Esrrb-/Tbx3-, Esrrb+/Tbx3-, Esrrb+/Tbx3+) (Hormoz et al. 2016). The gold standard target genes of Esrrb and Tbx3 was obtained by comparing Esrrb-/Tbx3-versus Esrrb+/Tbx3-samples and Esrrb+/Tbx3-versus Esrrb+/Tbx3+ samples with 2-fold change criterion, respectively. The target genes of Nanog and Pou5f1 were identified by using microarray data in mESC with Nanog and Pou5f1 knockdown (Loh et al. 2006). To identify target genes of these two TFs, we used a 2-fold change and a p-value < 0.01 as described in the original data analysis.

ChIP-seq data for Pou5f1, Esrrb, Nanog in mESCs were reanalyzed for peak calling (Chen et al. 2008). After removing the adapter sequence using CutAdapt (Martin 2011) implemented in TrimGalore-0.4.5, we aligned the ChIP-seq reads to the mm10 genome using Bowtie2 (Langmead et al. 2013). ChIP-seq peak was called against GFP control using the “findPeaks” command in the Homer package (Heinz et al. 2010).

### Robustness of the performance of TENET

In order to evaluate the robustness of TENET, we ran the Wishbone 57 times with different options on the Boolean expression data of single-cells obtained from early blood development experiments (Moignard et al. 2015). 57 Wishbone trajectories were obtained by running Wishbone with 19 different initial states provided in the reference paper (Moignard et al. 2015) and three different choices of cells based on the branches (total cells, trunk + first branch, trunk + second branch). The other GRN reconstructors from Beeline were also run based on these 57 Wishbone trajectories.

### Condition specific targets

To identify condition specific targets, we reconstructed GRNs using the pseudo-time ordered expression data of 2iL+NPCs and SL+NPCs using TENET. Subsequently, the condition specific targets of the top 20 factors in the common GRN were obtained by selecting targets in the culture condition specific GRNs. For example, the 35 target genes of Nme2 were included in the 2iL-specific but not in the SL-specific GRN whereas the 14 target genes were included in the SL-specific but not in the 2iL-specific GRN.

### ESC culture

E14 mESC were cultured on plastic plates coated with 0.1% gelatin (Sigma #G1393) in either DMEM knockout (Gibco #10829), 15% FBS (Gibco #10270), 1xPen-Strep-Glutamine (Gibco #10378), 1xMEM (Gibco #11140), 1xB-ME (Gibco #21985) and 1000U/mL LIF (Merck #ESG1107) (“Serum”) or in NDiff 227 (Takara #Y40002), 3uM CHIR99021 (Tocris #4423), 1uM PD0325901 (Tocris #4192) and 1000U/mL (“2iL”). For Nme2 experiments, mESCs were treated with either vehicle DMSO (Sigma #02660) or 0.5uM STP (StemCell technologies #72652) for 24 or 48hr.

### Alkaline phosphatase staining

For AP staining, 1000 mESCs were seeded in a 12-well plate and cultured for 24 or 48hr. The cells were washed in PBS, fixed in 1% formaldehyde and stained with AP following manufacturers instruction (Merck #SCR004). For quantification of positive stained colonies, four randomly selected areas of each well were imaged (10x magnification; Nikon Eclipse TS2) and manually counted. Colonies were marked as pluripotent or primed based on morphology and intensity of AP staining. The mean and standard error of mean (SEM) were calculated over four independent replicates.

### Cell proliferation assay

For proliferation assay, 100,000 mESCs were seeded in a 6-well plate in both 2iL and SL conditions. Cells were initially allowed to attach for 24hrs before treatment with either DMSO or STP. After either 24hr or 48hr of DMSO or STP treatment, cells were detached from the plate using Accutase and counted using the TC-20 automated cell counter (BioRad). Data are mean + SEM from 4 biological replicates.

### RNA extraction and qPCR

Total RNA was harvested using Trizol (Ambion #15596026), lock-gel columns (5prime #733-2478) and precipitated in chloroform/isopropanol using with glycogen. Reverse transcription was performed with 1ug of RNA using high capacity cDNA kit (Applied Biosystem #4368814). Quantitative-PCR was performed using SYBR-green with LightCycler480. To obtain relative gene expression levels, expression levels were normalized to Gapdh as a control.

## RESULTS

### TENET quantifies the strength of putative causal relationships between genes from scRNAseq data aligned along the pseudo-time

TENET measures TE for all pairs of genes to reconstruct a GRN. To assign time to the cells, TENET aligns cells along the pseudo-time. The paired gene expression levels along the pseudo-time are used to calculate TE (Figure 1a). Given the pseudo-time ordered expression profiles (Figure 1a), TE quantifies the strength of putative causal relationships of a gene *X* to a gene *Y* (Figure 1b) by considering the past events of the two genes. TE represents the level of information in gene *X* that contributes to the prediction of the current event *Y_t_*. The highly significant relationships between genes are obtained by modeling all possible relationships with normal distribution (Benjamini-Hochberg’s FDR (Benjamini and Hochberg 1995)<0.01). The potential indirect relationships are removed by applying data processing inequality measure (Margolin et al. 2006) (Figure 1c) (see Methods). TENET can be run on various sets of cell type regulators including either known set of genes or the set of all TFs, or even on the entire set of genes depending on the network of interest. After feature selection, the network analysis is applied to understand key regulators and relationships within the networks. In sum, TENET is useful in identifying target genes of a regulator and predicting key regulators.

### The TF target genes showed significantly higher TE values than randomly selected genes

We applied TENET to the scRNAseq data during mESC differentiation into NPCs (Tuck et al. 2018). We profiled mESCs cultured in 2iL (serum-free media with MEK and GSK3 inhibitors and cytokine LIF) and SL (serum media and cytokine LIF), and induced differentiation into neural progenitor cells (Bibel et al. 2007). The 2iL cultured mESCs (termed ‘ground state’) homogeneously express naïve pluripotency markers mimicking mouse epiblast, while SL cultures contain a heterogeneous mix of undifferentiated and differentiating ESCs (Alexandrova et al. 2016; Kalkan et al. 2017). Another motivation for choosing the mESC experimental system was that a number of ChIP-seq and RNAseq datasets are publicly available for validation (Chen et al. 2008; Loh et al. 2006; Hormoz et al. 2016). Visualization of the scRNAseq data during mESC differentiation using tSNE showed the differentiation trajectory from naïve ground state pluripotency (2iL) to differentiation-permissive (SL) to NPCs (Figure 2a). Consistent with the differentiation time course, general and naïve pluripotency markers including Pou5f1 (or Oct4), Sox2 and Nanog were highly expressed in the mESC population whereas NPC markers such as Pax6 and Slc1a3 were highly expressed in the NPCs (Figure S1). Then, we evaluated the TE values of the target genes supported by ChIP-seq at the promoter proximal (+/2kbps) region. We chose c-Myc, n-Myc, E2f1 and Zfx (Chen et al. 2008) as their occupancy is often observed at the promoter region of their target genes. Applying peak calling using Homer (Heinz et al. 2010), we found 541 c-Myc promoter proximal peaks. The TE values of the c-Myc targets were compared to the randomly selected genes (as control) with the same sample size, similar GC contents and expression levels. Repeating the process 1,000 times, we observed that the 541 c-Myc target genes showed significantly higher TE values (p-value=1.19e-27) than the randomly selected genes (Figure 2b). We also confirmed that ChIP-seq binding targets for other promoter binding TFs such as n-Myc, E2f1 and Zfx also have significantly higher TE values compared with the random targets (Figure S2a-c).

**Figure 2.**
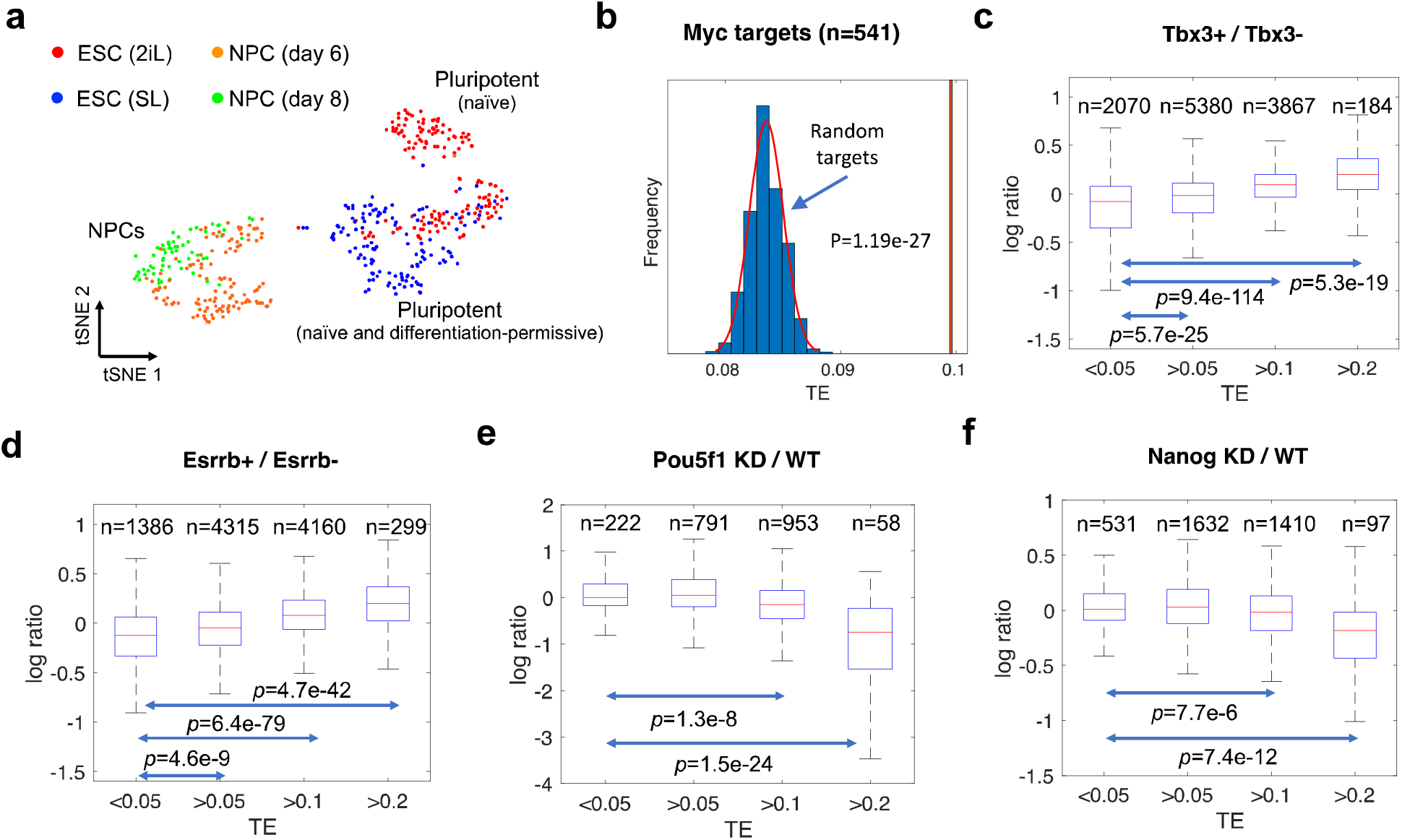
Validation of TENET-inferred GRNs for the mouse embryonic stem cell (mESC) pluripotency. **a**. A tSNE plot of the mESCs (2iL and SL) and NPCs shows distinct expression. **b**. The c-Myc target genes have higher TE values than the randomly selected 541 genes (repeated 1000 times). The expression ratio of predicted Tbx3 (**c**) or Esrrb (**d**) target genes (Tbx3 or Esrrb overexpression (Tbx3+ or Esrrb+) against control (Tbx3− or Esrrb−)). The expression ratio of predicted Pou5f1 (**e**) or Nanog (**f**) target genes (knockdown versus wild-type).

Additionally, we performed evaluation of TE values using the scRNAseq dataset for the reprogramming of mouse fibroblasts into induced cardiomyocytes (Liu et al. 2017b). Investigation using Gata4 ChIP-seq in cardiomyocytes (Luna-Zurita et al. 2016) confirmed that the 331 potential target genes with Gata4 promoter occupancy also possess significantly higher TE values compared with random targets (Figure S2d).

### TE values reflect the degree of dependency to the regulator

Gene perturbation followed by gene expression measurement by bulk RNAseq has been widely used to determine potential target genes. We further examined the TE values of the potential TF target genes identified by overexpression of Esrrb and Tbx3 as well as knockdown of Pou5f1 and Nanog (Loh et al. 2006; Hormoz et al. 2016). We divided the genes based on their TE values and investigated the fold changes upon the perturbation of the corresponding TF. Interestingly, the expression levels of the genes with low TE values (<0.05) had little or no influence upon perturbation. However, the expression levels of the genes with high TE values were markedly increased upon overexpression of Esrrb and Tbx3 and consistently, decreased upon knockdown of Pou5f1 and Nanog. These changes were particularly more significant for genes with higher TE values (>0.2) (Figure 2c-f). These results indicate that TE values reflects the degree of dependency of the target genes to the expression of their regulator.

### TENET can predict key regulators from scRNAseq data

To determine whether the TENET captures the key biological processes, we investigated hub nodes and evaluated if key regulators were well represented. From the reconstructed GRNs from mESC to neural cells, we assessed if key regulators (based on the number of outgoing edges) in the GRNs are associated with stem cell or neural cell biology. The gene ontology (GO) terms and KEGG pathways enrichment tests showed that the hub regulators (number of outgoing edges >= 5) are mostly associated with pluripotent stem cells and cellular differentiation functions (Figure S3a). We also benchmarked and compared TENET’s performance to other methods including SCODE (Matsumoto et al. 2017), GENIE3 (Huynh-Thu et al. 2010), GRNBOOST2 (Moerman et al. 2019), SINCERITIES (Papili Gao et al. 2018), LEAP (Specht and Li 2017), SCRIBE (Qiu et al. 2020), and SCINGE (Deshpande et al. 2019). For unbiased comparison, we ran each GRN method on the same set of 3,277 highly variable genes (see Methods).

The top 4 ranked regulators determined by TENET were markers for pluripotency (Pou5f1, Nanog, Esrrb, and Tbx3) (Figure 3a). Compared to TENET, most methods failed to identify these key genes in the hub list except for SCRIBE. For instance, GENIE3 and GRNBOOST2 only found Nanog as the 14th and 5th of the top regulators, respectively; but they did not detect Pou5f1. SCRIBE, another TE-based GRN predictor identified Nanog, Pou5f1, Esrrb, Tbx3 as the top regulators, suggesting the algorithmic advantages of TE especially for scRNAseq data (Figure S4). However, both SCRIBE and SCODE the numbers of target genes in the hub node were drastically reduced beyond the 5th regulator, which highlights that these methods emphasize on a few potential regulators during network reconstruction.

**Figure 3.**
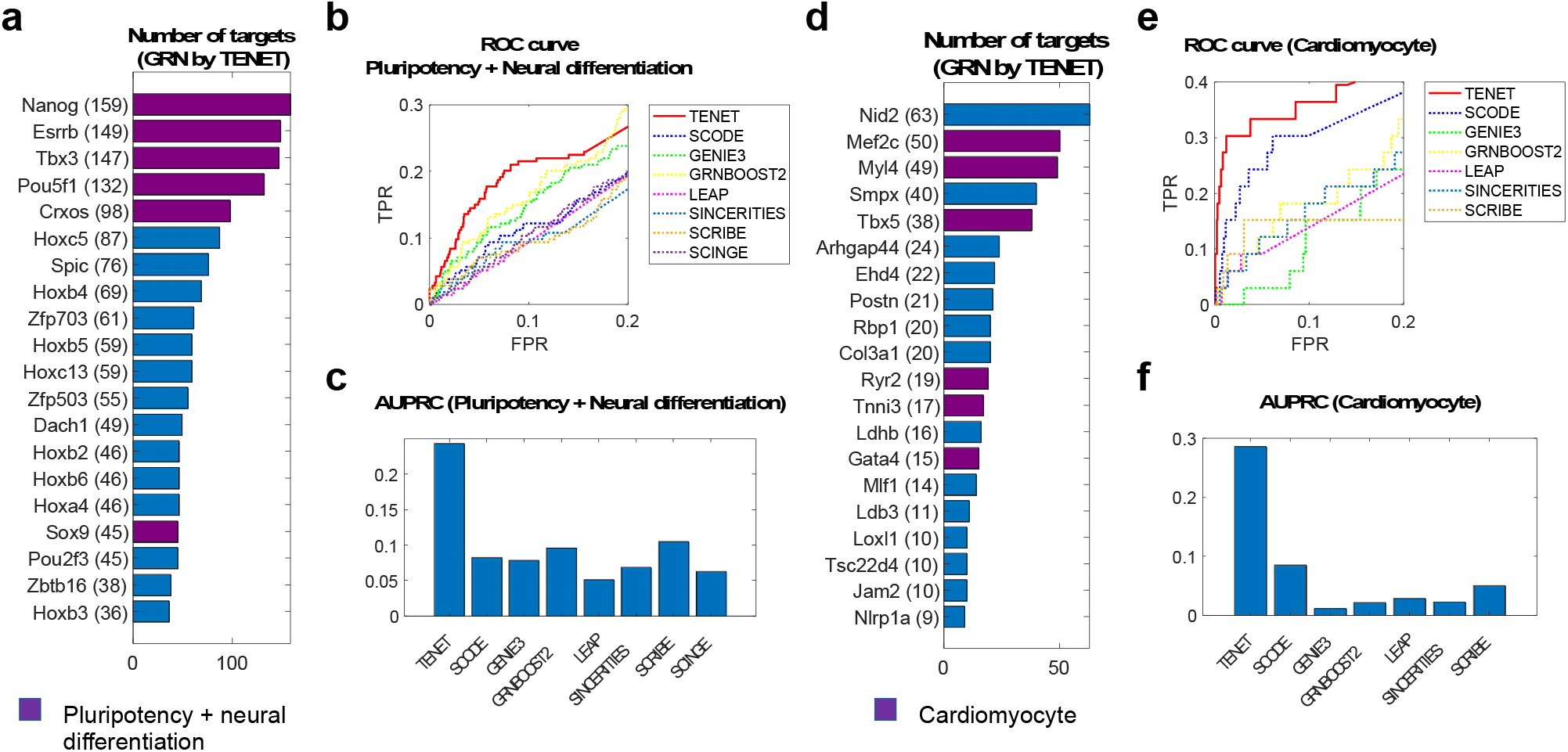
TENET outperformed other tools when predicting key regulatory factors for mESC pluripotency and direct reprogramming from mouse fibroblast into cardiomyocyte. **a**. Key regulatory factors for mESC pluripotency and neural differentiation predicted by TENET. The purple bar denotes pluripotency and neural differentiation genes. **b**. ROC curves and **c**. Area Under Precision-Recall Curves (AUPRCs) for the prediction of key regulatory factors of pluripotency and neural differentiation. **d**. Key regulatory factors for direct reprogramming into cardiomyocyte in the TENET-inferred GRN. Three major reprogramming factors Mef2c, Tbx5, Gata4 have a large number of targets. **e**. ROC curves and **f**. AUPRCs for the prediction of key regulatory factors of cardiomyocyte.

Intrigued by this, we investigated whether the hubs in the networks are associated with “pluripotency” or “neural differentiation” using the list of the genes obtained from GO database (see Methods). We investigated both receiver operating characteristic (ROC) curves and precision-recall curves (PRCs) while regarding genes in the GO database as true. The ROC curves for the pluripotency and neural differentiation demonstrates TENET’s far exceeding capability in predicting key regulatory factors related with these key GO terms compared to other methods (Figure 3b and S5a-b). The area under precision-recall curve (AUPRC) further confirmed the increased performance of TENET in capturing key regulators (Figure 3c). As TE values rely on pseudo-time, we also investigated whether TENET results were sensitive to other pseudo-time inference methods. Computing pseudo-time using PAGA (Wolf et al. 2019) and Slingshot (Street et al. 2018) showed that TENET is robust to the choice of pseudo-time inference and outperformed other GRN reconstructors (Figure S6).

To further test if TENET can suggest key regulatory factors in various biological systems, we reconstructed a GRN based on the scRNAseq data for direct reprogramming of mouse fibroblast into cardiomyocyte by overexpressing Mef2c, Tbx5 and Gata4 (Liu et al. 2017b). We first examined if these overexpressed factors were well predicted in the inferred GRNs. Consistently, TENET identified those three major reprogramming factors (Mef2c, Tbx5 and Gata4) as well as other genes associated with cardiomyocytes as top ranked regulators (Figure 3d and Figure S3b). Not surprisingly, these factors were not well observed in the GRNs inferred by other reconstruction methods (Figure 3e-f and S5c-d) with exception of SCRIBE that only found Mef2c (Figure S7). Additionally, while GRNBOOST2 showed relatively better performance in detecting pluripotency and neuronal differentiation factors, it failed in detecting the key factors during cardiomyocyte reprogramming. Collectively, our results show that TENET can robustly capture key regulatory genes for biological processes.

### TENET’s hub nodes were associated with the controllability of Boolean network dynamics

To further investigate the characteristics of TENET in finding key regulators, we compared the reconstructed networks with Boolean networks (BNs) (Moignard et al. 2015). BNs consider all possible binary status of its members (genes) and have been widely used to model biological systems (Li et al. 2004; Choi et al. 2012; Wang et al. 2010). BNs can simulate overexpression or knock-out of a gene and its consequences from the inferred networks. Therefore, BNs can be used to evaluate how much a member (i.e. gene) can influence the steady-state dynamics of the networks, and in combination with other members (called “controllability”) (Kim et al. 2013). Previously, a BN based GRN using 20 TFs was built using single cell qRT-PCR during mouse early blood development (Moignard et al. 2015) (Figure S8). Using the BN-inferred GRN as a surrogate for the gold standard, we first evaluate if the networks from GRN reconstructors accurately mimic the BN-inferred GRN. The comparison showed that TENET and GRNBOOST2 outperforms other approaches in both directed and undirected networks in this example (Figure S9a-b).

In the BNs, the number of final stable states (known as attractors) can be calculated while simulating all possible states of the members except one member of interest (a gene with perturbation). A critical member usually has a small number of attractors. Therefore, the predicted hub genes in the GRN will negatively correlate with the number of attractors if the hub genes are the key genes. In a series of experiments, TENET showed an ability to find key regulators. We further tested if the predicted key regulators are negatively correlated with the attractors found in the BNs. Our simulation showed that the TENET-inferred network has the strongest negative correlation with the number of attractors compared followed by SCRIBE, SCODE and GRNBOOST2 (Figure S9c), while other methods showed either no or positive correlation. This further demonstrated that TENET has the capability to identify key regulators.

### TENET outperforms other GRN reconstruction algorithms in identifying target genes

To further assess TENET, we used Beeline (Pratapa et al. 2020), a benchmarking software for GRN inference algorithms. Among them, we performed benchmarking only for those algorithms that can implement large scale GRN reconstruction including SCODE (Matsumoto et al. 2017), GENIE3 (Huynh-Thu et al. 2010), GRNBOOST2 (Moerman et al. 2019), SINCERITIES (Papili Gao et al. 2018), LEAP (Specht and Li 2017), SCRIBE (Qiu et al. 2020), and SCINGE (Deshpande et al. 2019), using the mESC scRNAseq dataset (Tuck et al. 2018). To prepare stringent datasets for evaluation, we regarded a target as true if the expression of the target gene is changed significantly by the perturbation study (Loh et al. 2006; Hormoz et al. 2016) and the binding occupancy of the corresponding regulator is observed nearby (+/-50kbps to transcription start sites (TSSs)) (Chen et al. 2008) (see Figure S10a and Methods).

We benchmarked all methods running Beeline on 3,277 highly variable genes (see Methods). Beeline (Pratapa et al. 2020) provided comprehensive results after running all GRN reconstructors. The ROC curves showed that TENET, GENIE3 and LEAP outperformed other reconstructors in predicting targets of Nanog, Pou5f1, Esrrb, and Tbx3 (Figure S10b-c). Interestingly, SCRIBE showed worse performance than TENET, while GENIE3 and LEAP failed to find key regulatory genes but showed good performance in this benchmarking test (even with a small number of regulators).

### TENET identifies culture condition specific regulators

To search for potential regulators besides the known TFs during stem cell differentiation (Tuck et al. 2018), we extended the GRN by considering 13,694 highly variable genes as well as target genes (see Methods). In addition to several known pluripotency (Nanog, Sox2, Pou5f1, Tfcp2l1 etc.,) and neural regulators (Meis1, Tbx3 etc.,), we were intrigued to find Fgf4 and Nme2 as the top regulators (ranked by number of targets, Figure S11b and Table S1). Fgf4 is known to be dispensable for embryonic stem cells, but is critical for exit from self-renewal and differentiation (Almousailleakh et al. 2007; Lanner and Rossant 2010), while Nme2 (Zhu et al. 2009) has been implicated in stem cell pluripotency (Figure S11).

As we profiled mESCs in both ground-state 2iL and heterogeneous SL conditions, we assessed whether TENET could further distinguish them and identify culture-condition specific GRNs. We reconstructed GRNs for 2iL and SL condition separately and compared the regulators as well as their specific targets (Figure S12, see Methods). We found several naïve pluripotency markers specifically enriched in 2iL condition including Nanog (Wray et al. 2010), Esrrb (Martello et al. 2012), and Tfcp2l1 (Martello et al. 2013; Qiu et al. 2015), whereas heterogeneous and hypermethylated SL condition (Habibi et al. 2013; Leitch et al. 2013) regulators included Tet1 (Pantier et al. 2019; Ito et al. 2010) and Dnmt3l (Ficz et al. 2013) and Zfp57 (Riso et al. 2016) (Figure S12). Interestingly, Nme2 was a top mESC regulator for the 2iL GRNs. Intrigued by this, we sought to investigate the effect of Nme2 perturbation using an small molecule inhibitor Stauprimide (STP) that blocks the nuclear localization (Zhu et al. 2009). We cultured mESCs in 2iL and SL conditions and treated cells with 0.5μM STP (Figure 4a). The cellular proliferation and division were significantly inhibited in both culture conditions, but were drastic in 2iL (Figure 4b). Upon 24hr STP treatment in 2iL, we could visually observe high levels of apoptotic cells, detached colonies and few viable cells at 48hrs. We quantified the STP effect on pluripotency by alkaline phosphatase staining (AP; Figure 4c) and scored cells either as *undifferentiated* (high AP staining, rounded colonies; naïve mESCs) or as *mixed* (low/no AP staining, flattened colonies; differentiation-like/apoptotic cells) (Kalkan et al. 2017; Liu et al. 2017a). The STP treatment in 2iL led to a drastic decrease in undifferentiated colonies (45% ± 5.9% colonies) and the remaining mixed cells were mostly composed of apoptotic cells.

**Figure 4.**
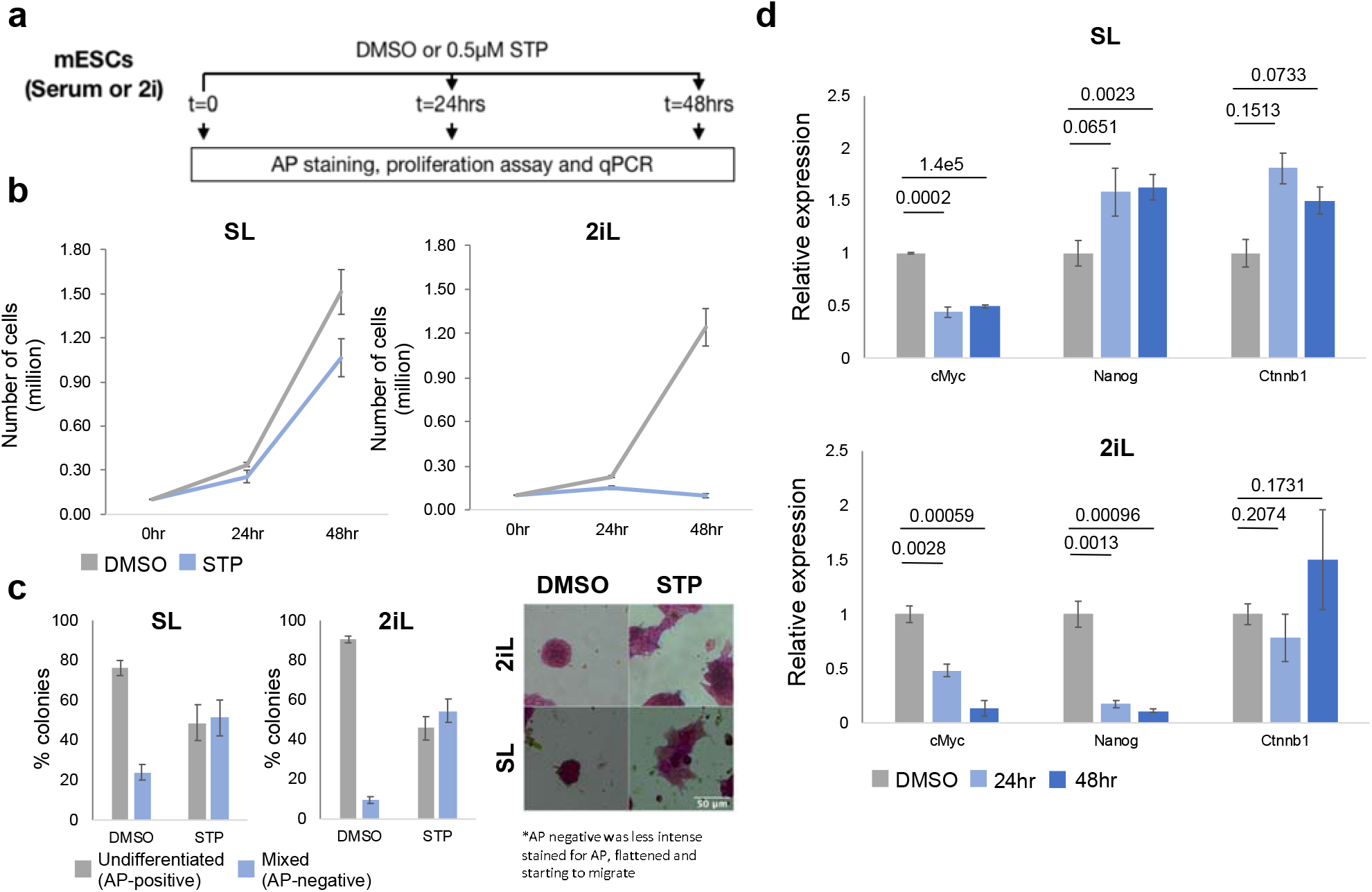
Nme2 inhibition blocks proliferation of mESC in 2iL condition. **a**. Experimental design: The mESCs were seeded in either SL or 2iL culture conditions and were treated with either DMSO (control) or 0.5μM STP for 24hr and 48hr. The 6 samples were assayed for proliferation rates, relative transcript expression and for pluripotency using alkaline phosphatase (AP). **b**. Cell proliferation assay for mESCs cultured in SL and 2iL conditions with either DMSO (Control) or 0.5μM STP. The mean and SEM were calculated over four independent replicates. **c**. STP treatment leads reduction in undifferentiated colonies and increase in mixed differentiation-like colonies across both SL (48hrs) and 2iL (24hrs) conditions, based on AP staining. AP staining intensity and colony morphology are used to classify undifferentiated (high AP staining, rounded colonies) and mixed (low/no AP staining, flattened colonies; differentiation-like; apoptotic cells) populations. Representative images of undifferentiated and mixed colonies in control and STP treated colonies across both culture conditions. The data in the barplots describe the mean ± SEM from 2 biologically independent replicates. **d**. The c-Myc transcript levels are down regulated both in 2iL and SL upon STP treatment, owing to impaired Nme2 nuclear localization. The Nme2 target genes in TENET (Nanog and Ctnnb1) are selectively regulated between culture conditions. Significance (p-value) are highlighted above barplots. The data in barplots describe the mean ± SEM from 3 biologically independent replicates.

Previously, c-Myc has been reported as a target gene of Nme2 (Zhu et al. 2009), consistent with TENET prediction. We confirmed that c-Myc expression was significantly downregulated upon STP treatment in both culture conditions (Figure 4d). TENET also predicted several TFs including Nanog and Ctnnb1 at the target of Nme2. We found that both Nanog and Ctnnb1 transcripts were highly upregulated upon STP treatment in both culture conditions but more significant in 2iL condition, indicating condition specific regulation of Nme2 as predicted by TENET (Figure 4d).

## DISCUSSION

Systems biology approaches to infer GRNs can provide a hypothesis for further experimental validation. Existing methods for bulk transcriptomics datasets are limited because they cannot capture the continuous cellular dynamics and/or require cell synchronization to avoid “average out” expression. scRNAseq has emerged as an alternative because of its power to provide the transcriptomic snapshots of hundreds, thousands of cells on a massive scale, from same population. Subsequently, computational approaches used scRNAseq for GRN reconstruction (Moignard et al. 2015; Sanchez-Castillo et al. 2018; Matsumoto et al. 2017; Moerman et al. 2019; Papili Gao et al. 2018; Specht and Li 2017; Deshpande et al. 2019; Qiu et al. 2020; Møller and Natarajan 2020; Aibar et al. 2017).

Many GRN reconstruction algorithms including TENET use the temporal gene expression changes, after ordering cells across pseudo-time. For example, GENIE3 (Huynh-Thu et al. 2010) and GRNBOOST2 (Moerman et al. 2019) were originally applied the ensembles of regression trees to temporal bulk expression data. LEAP (Specht and Li 2017) calculates possible maximum time-lagged correlations. SINCERITIES (Papili Gao et al. 2018) and SCINGE (Deshpande et al. 2019) used Granger causality from pseudo-time ordered data. SCODE (Matsumoto et al. 2017) uses a mechanistical model of ordinary differential equations on the pseudo-time aligned scRNAseq data. Compared with current methods, TENET makes use of the power of information theory by adopting TE on gene expression along the pseudo-time. Therefore, the performance of these predicted regulators could be dependent on the performance of the pseudo-time inference. However, we found that TENET is robust to the multiple pseudo-time inference approaches in comparison with other GRN reconstructors (Figure S6).

We showed that TE values of the known target genes were significantly higher than randomly selected genes (Figure 2b and S2). The target genes with higher TE values were more significantly perturbed by either overexpression or knockdown of the corresponding regulators (Figure 2c-f). We also performed comprehensive benchmarking of TENET and several GRN reconstructors using Beeline (Pratapa et al. 2020) and its automated pipeline. TENET was consistently one of the top performing GRN reconstructors in these tests.

The evaluation of the performances of GRN reconstructors by counting the number of true or false prediction does not fully reflect the importance of the inferred network. We observe that TENET consistently predicts and identifies key regulators. This is important because upstream regulators for a biological process are often of interest to explain the underlying mechanisms. It is still required to evaluate if the inferred networks reflect the key underlying biological processes. Applying TENET to a series of scRNAseq datasets including 1) mESC differentiation and 2) reprogramming to cardiomyocytes, we find that TENET identified key factors as the top scoring hubs. For mESC differentiation, TENET ranked Nanog, Pou5f1, Esrrb and Tbx3 as the top 4 regulators, while existing methods failed to identify these key factors. In an additional test using GO terms, TENET identified gene relationships associated with pluripotency and neural differentiation (Figure 3b-c). Interestingly, existing methods including LEAP and SINCERITIES did not find any genes related to pluripotency in their networks (Figure S5b). Analyzing the reprogramming to cardiomyocytes scRNAseq data, only TENET identified the reprogramming factors (Mef2c, Tbx5 and Gata4) (Liu et al. 2017b) (Figure 3d-f and Figure S7). These results suggest that while other approaches successful in finding some regulatory rules, they cannot make networks focusing on the key biological process.

We further questioned if TENET is capable of identifying key regulators using BNs. While BNs may not be a perfect model of biological system, they can still provide a comprehensive systematic overview by visiting all potential states. In BN, the key nodes usually have small number of attractors as they drive the networks into more determined status. In our analysis using BNs, TENET-inferred networks were negatively correlated with the number of attractors (Figure S9), indicating the key ability to capture biological processes.

A number of studies showed distinct expression patterns in the pseudo-space (Halpern et al. 2018; Nowotschin et al. 2019). Since pseudo-time inference can lead to multiple branched trajectories, we also applied TENET to individual branches. These expression changes for some genes may be attributed to association along the spatial axis. However, the associating potential causal relationships for them may not be relevant.

With the power to predict key regulators, we applied TENET to identify mESC culture-condition specific regulators. TENET predicted several TFs (Nanog, Esrrb, and Nme2) as specific for 2iL compared to SL culture conditions (Figure S12). Although Nme2 is expressed both in 2iL and SL, perturbing Nme2 leads to more dramatic effects (reduced proliferation, AP staining and apoptosis) in the 2iL condition; consistent with our prediction. In sum, TENET is a useful approach to predict previously uncharacterized regulatory mechanisms from scRNAseq.

## DATA AVAILABILITY

A source code for TENET and input files for the benchmarking datasets are available at https://github.com/neocaleb/TENET.

## ACKNOLEDGEMENT

We are grateful to Patrick Martin for proofreading of the manuscript and Sen Li for updating TENET.

## AUTHOR CONTRIBUTIONS

K.J.W. conceived and designed TENET. J.K., S.T.J. and K.N.N. performed the experiments and analyzed the data. J.K., K.J.W. and K.N.N. wrote the paper.

