## Supplementary Material for "TENET: Gene network reconstruction using transfer entropy reveals key regulatory factors from single cell transcriptomic data"

[1]Biotech Research and Innovation Centre (BRIC), University of Copenhagen, 2200 Copenhagen N, Denmark, [2]Novo Nordisk Foundation Center for Stem Cell Biology, DanStem, Faculty of Health and Medical Sciences, University of Copenhagen, Ole Maaløes Vej 5, 2200 Copenhagen N, Denmark, [3]Functional Genomics and Metabolism Unit, Department of Biochemistry and Molecular Biology, University of Southern Denmark, Denmark, [4]Danish Institute of Advanced Study (D-IAS), University of Southern Denmark, Denmark

### Supplementary Materials

* To whom correspondence should be addressed to KJW (Tel: +45-353-31419 ; Fax: +45-726-20285). Correspondence may also be addressed to KNN (Tel: +45-655-08929;).

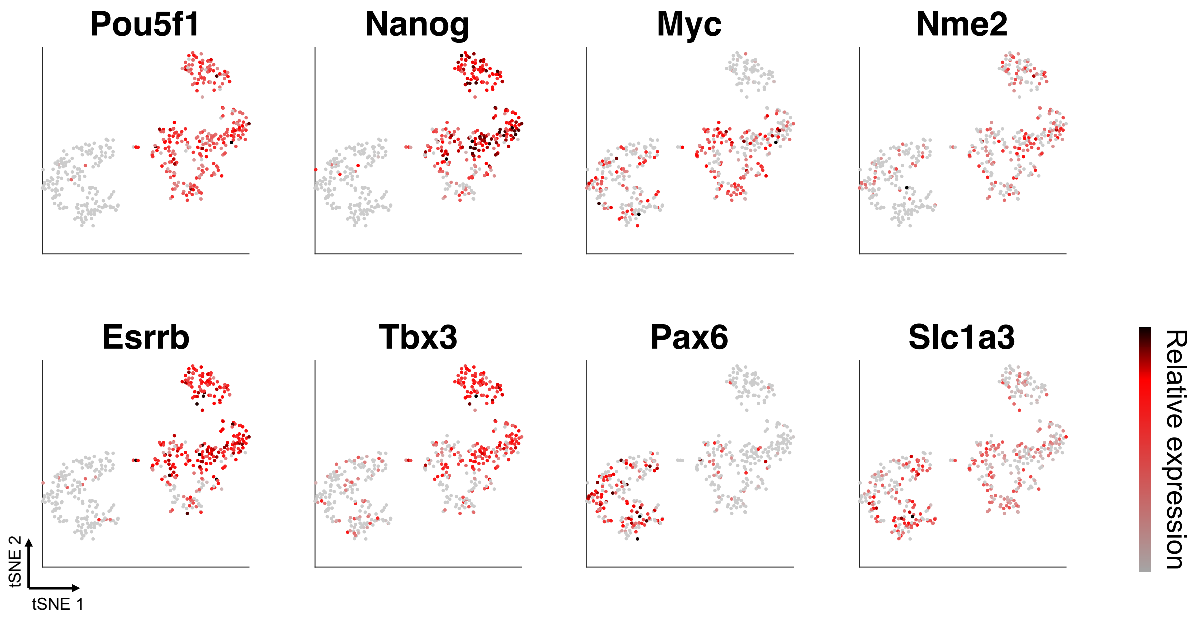


**Figure S1. Relative expressions of eight marker genes on tSNE plots of the mESC (2i and serum) and NPCs**

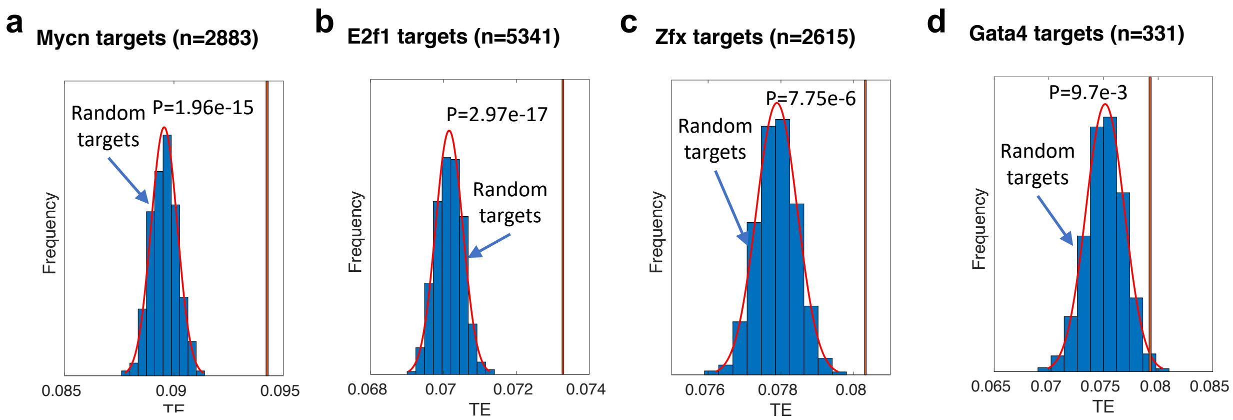


**Figure S2. TENET-inferred targets are confirmed by binding occupancy at the promoter regions in mouse embryonic stem cells** (1) **and induced cardiomyocytes** (2)**.** The target genes of n-Myc (**a**), E2f1 (**b**), and Zfx (**c**) in mouse embryonic stem cells and Gata4 (**d**) in induced cardiomyocytes have significantly higher TE values than random sets. The red graph represents a fitted normal distribution obtained from 1000 random sets of genes.

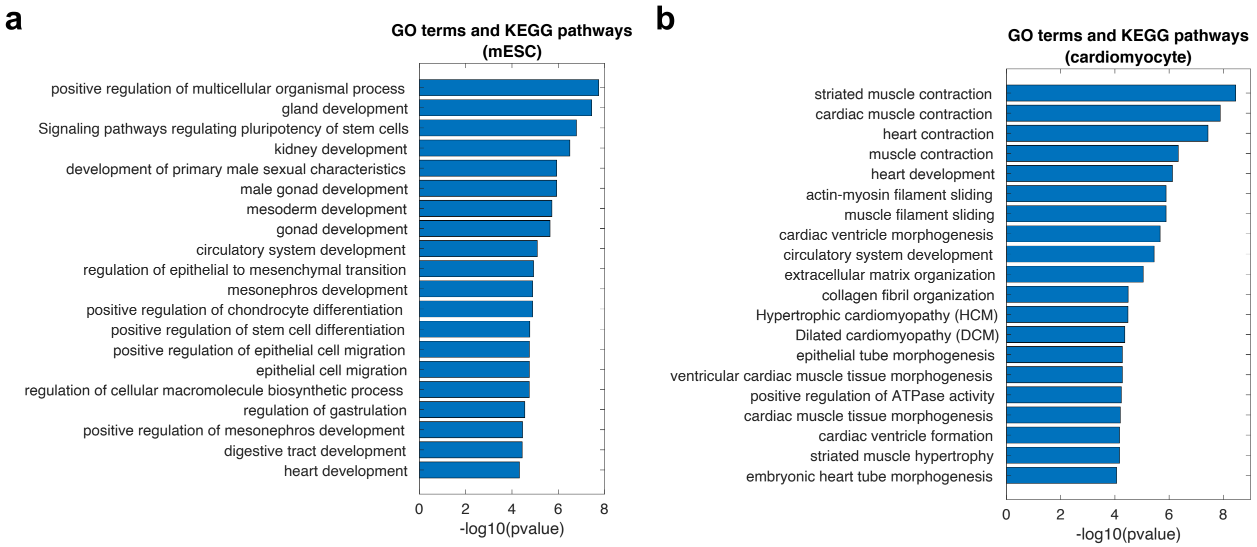


**Figure S3. The GO terms and KEGG pathways associated with the hub regulators (number of outgoing edges >=5) in mouse embryonic stem cells (a) and induced cardiomyocyte (b).**

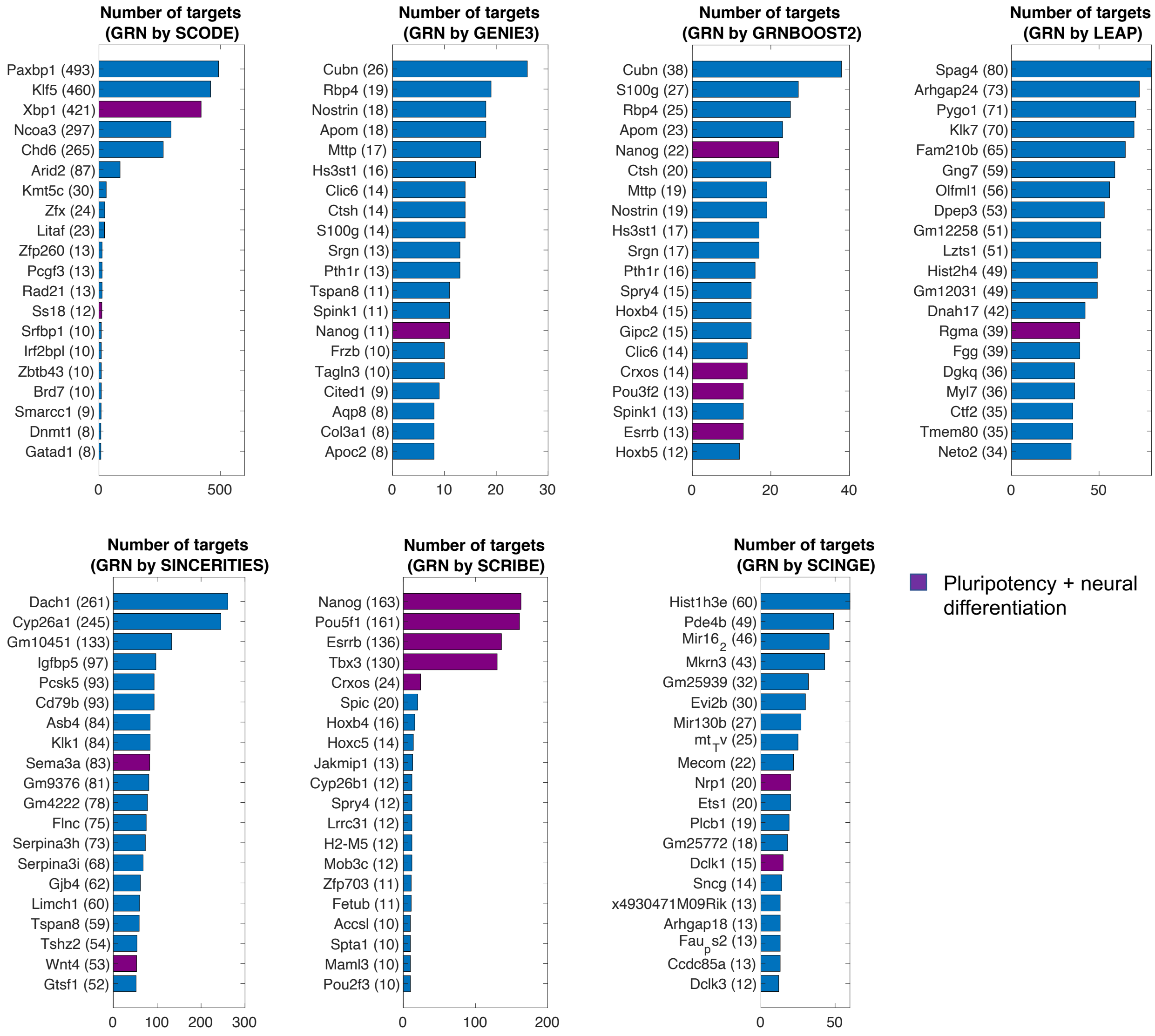


**Figure S4. Key regulators for mESC pluripotency and neural differentiation predicted by seven GRN reconstruction algorithms.**

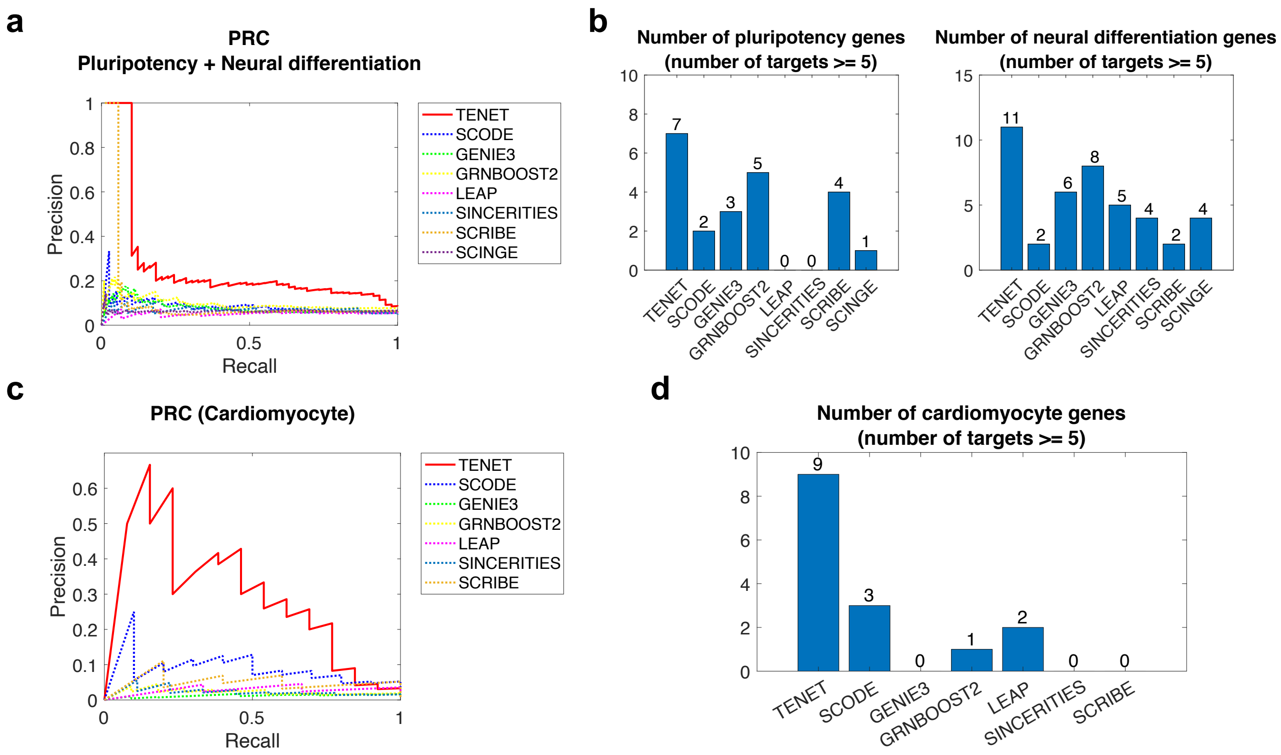


**Figure S5. TENET identified key regulatory factors for mESC pluripotency and direct reprogramming from mouse fibroblast into cardiomyocyte** (2)**. a.** Precision-Recall Curves (PRCs) for predicting key regulatory factors of pluripotency and neural differentiation. **b**. The number of hub genes (number of targets >= 5) related with pluripotency genes and neural differentiation genes. **c.** PRCs for predicting key regulatory factors of cardiomyocyte. **d**. The number of hub genes (number of targets >= 5) related with genes related to cardiomyocyte.

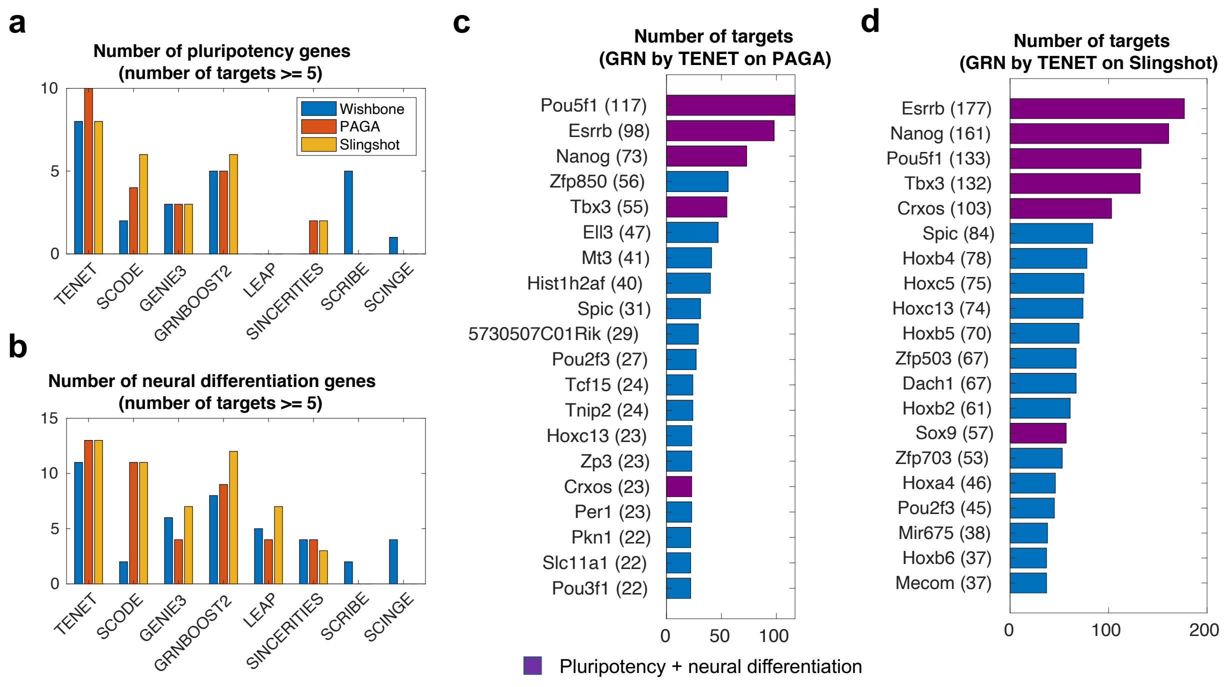


**Figure S6. TENET outperformed in terms of predicting key regulator factors for mESC pluripotency regardless of pseudo-time inference methods.** The number of hub genes (number of targets >= 5) related with pluripotency genes (**a**) and neural differentiation genes (**b**). Key regulatory factors for mESC pluripotency and neural differentiation predicted by TENET based on PAGA (**c**) and Slingshot (**d**)

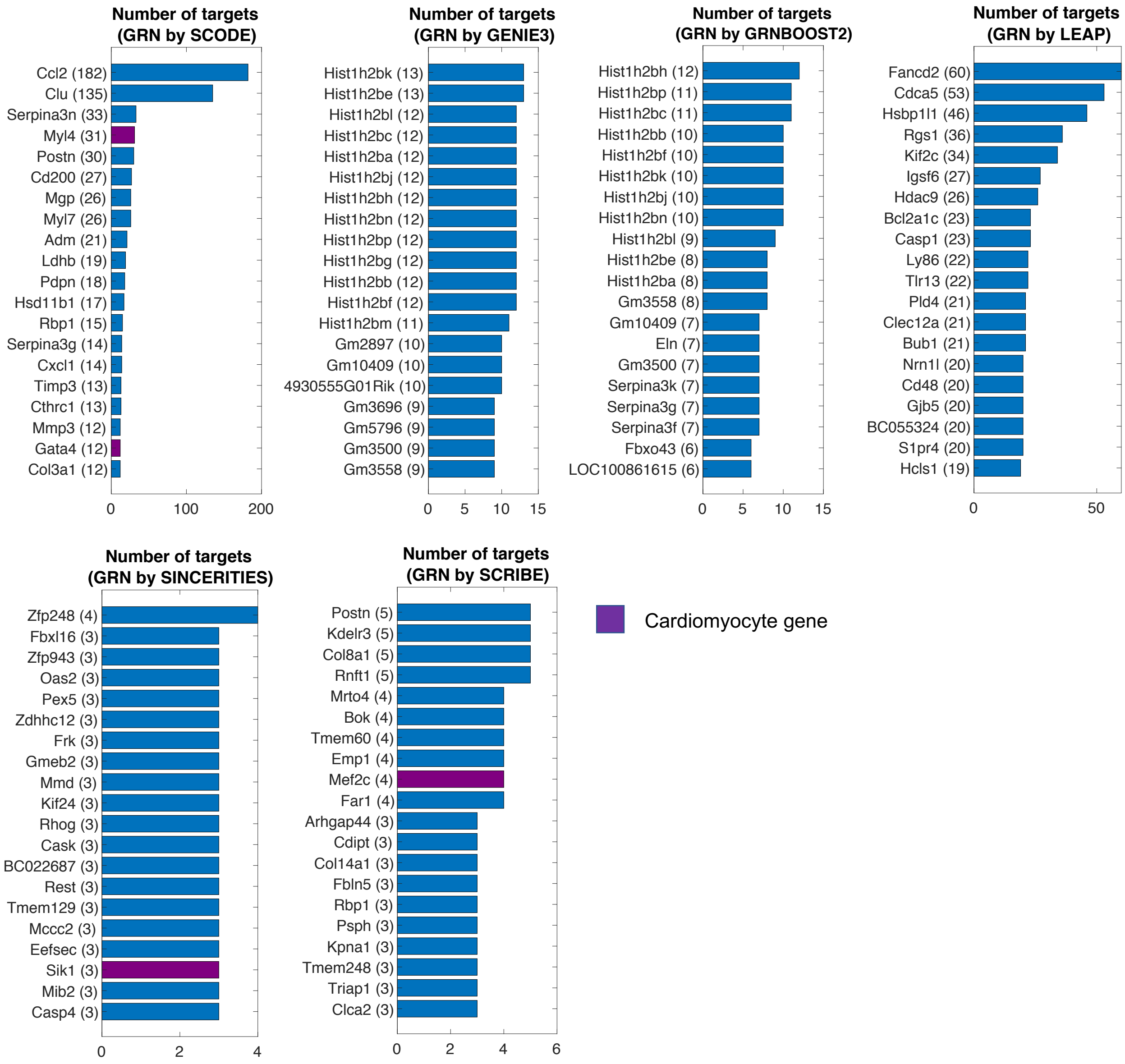


**Figure S7. Key regulatory factors for cardiomyocyte direct reprogramming predicted by six GRN inference algorithms.**

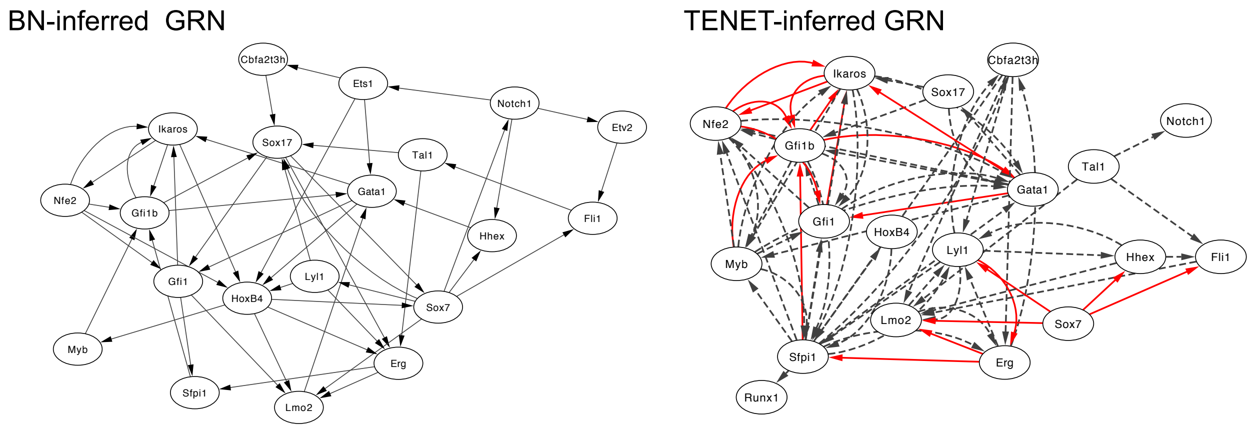


**Figure S8**. **BN (Boolean network)** (3)**-inferred GRN (left) and TENET-inferred GRN (right) for early blood development.** The red solid links indicate the correctly identified links by TENET. The black dashed links denote false positive.

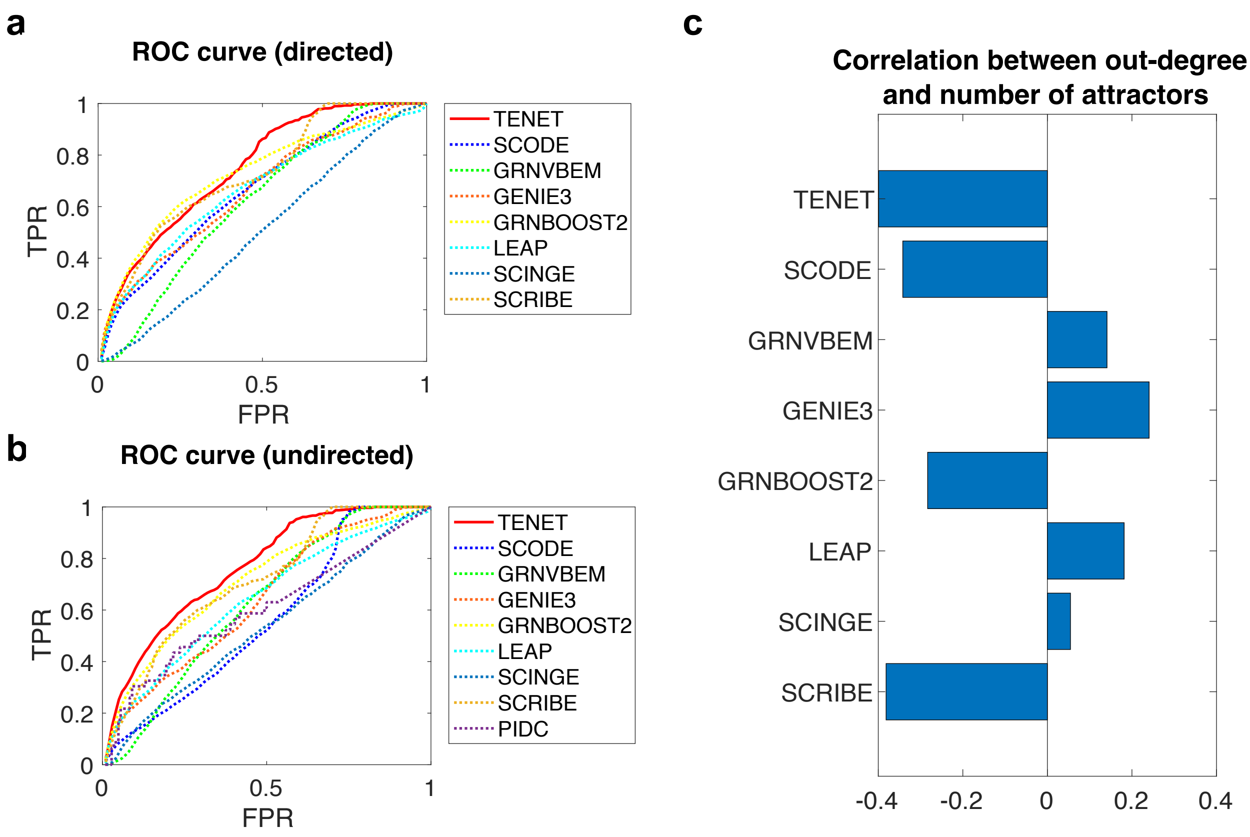


**Figure S9**. **TENET mimics the controllability of a Boolean network model for mouse early blood development** (3). **a**. Average ROC curves for the GRN of early blood development by TENET and seven different algorithms. Average of ROC curve for each algorithm was obtained from 57 GRNs based on different Wishbone results with different options (see Methods). **b**. Average ROC curves of TENET and eight different algorithms considering undirected links. We merged incoming and outgoing links into one link. **c**. Correlation between the out-degree and the number of attractors.

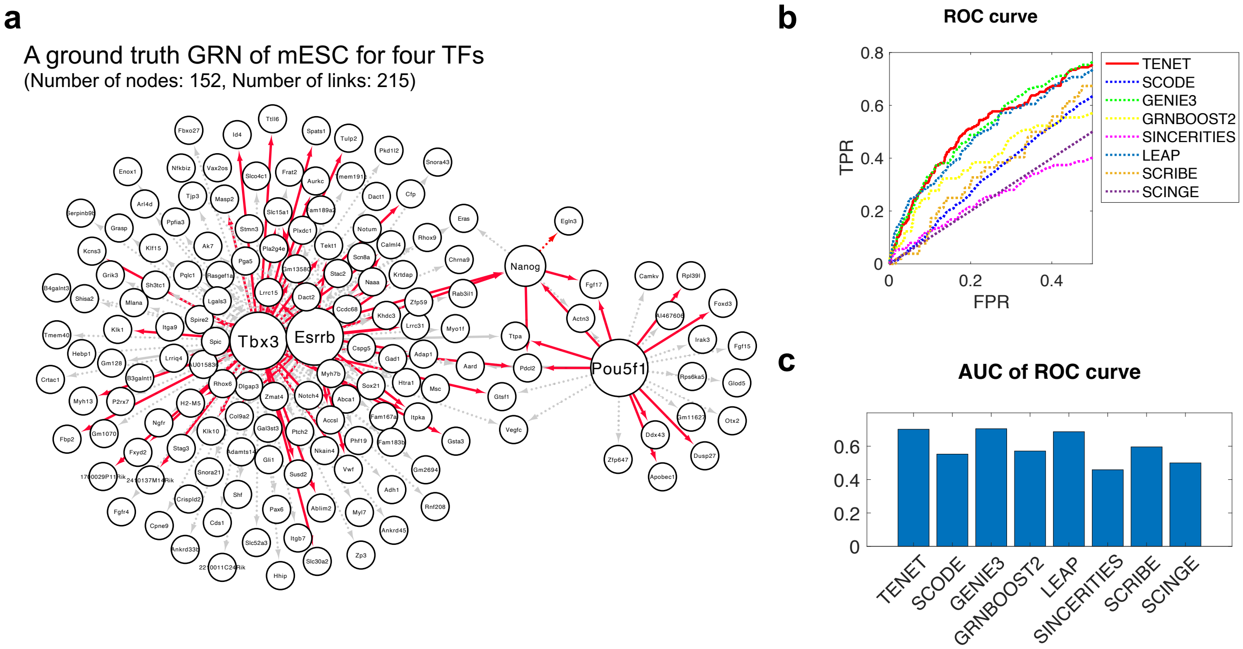


**Figure S10. The performance of TENET on reconstruction of GRN for the mESC pluripotency. a.** A ground truth GRN of mESC for four pluripotency TFs (Nanog, Pou5f1, Esrrb, and Tbx3). The red solid links indicate the correctly identified links by TENET. The grey dashed links denote false positive. **b**. ROC curves for the mESC GRN by TENET and seven different algorithms. **c**. Comparison of area under curve (AUC).

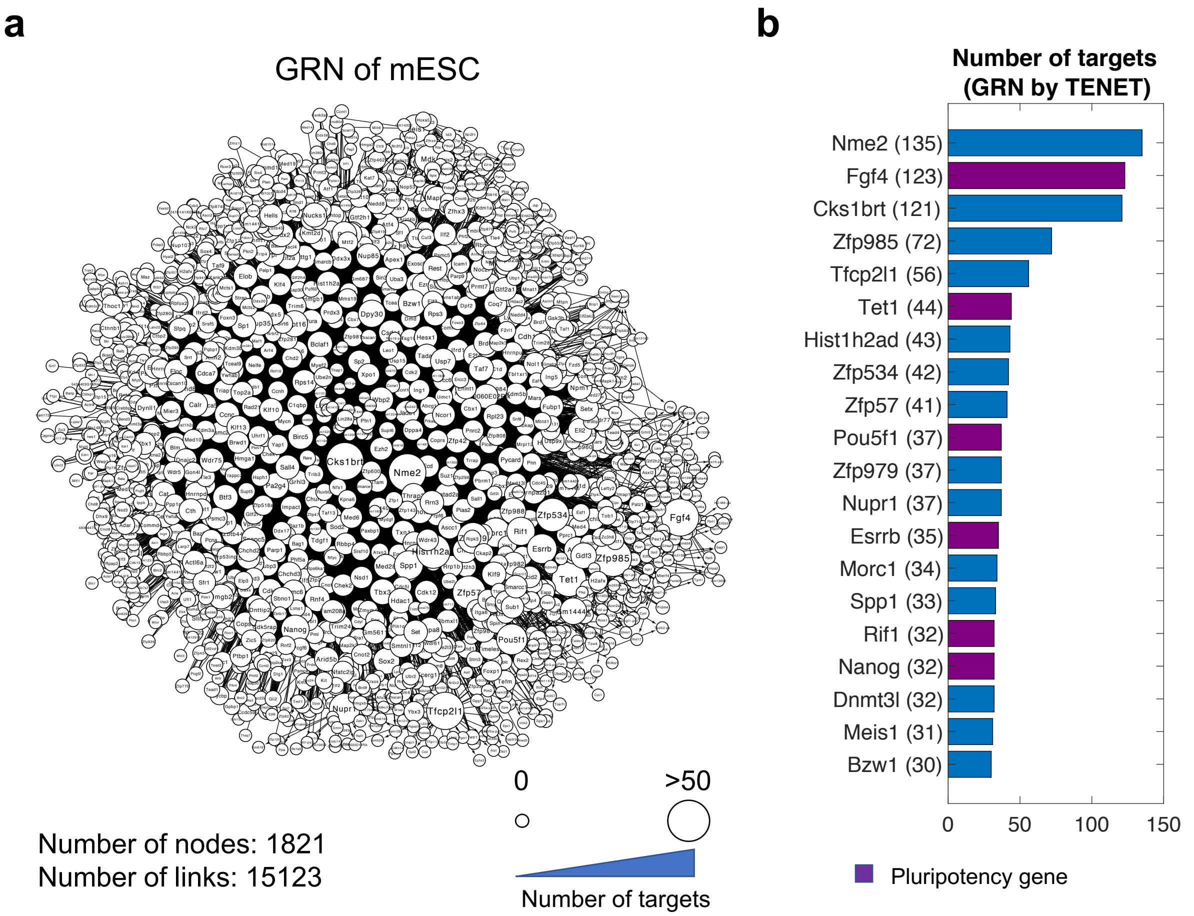


**Figure S11**. **TENET identified a critical factor Nme2 as the top scoring factor in terms of number of targets. a**. TENET-inferred GRN of mESC. **b**. Number of targets for 20 top TFs in the TENET-inferred GRN. Nme2 has the largest number of targets.

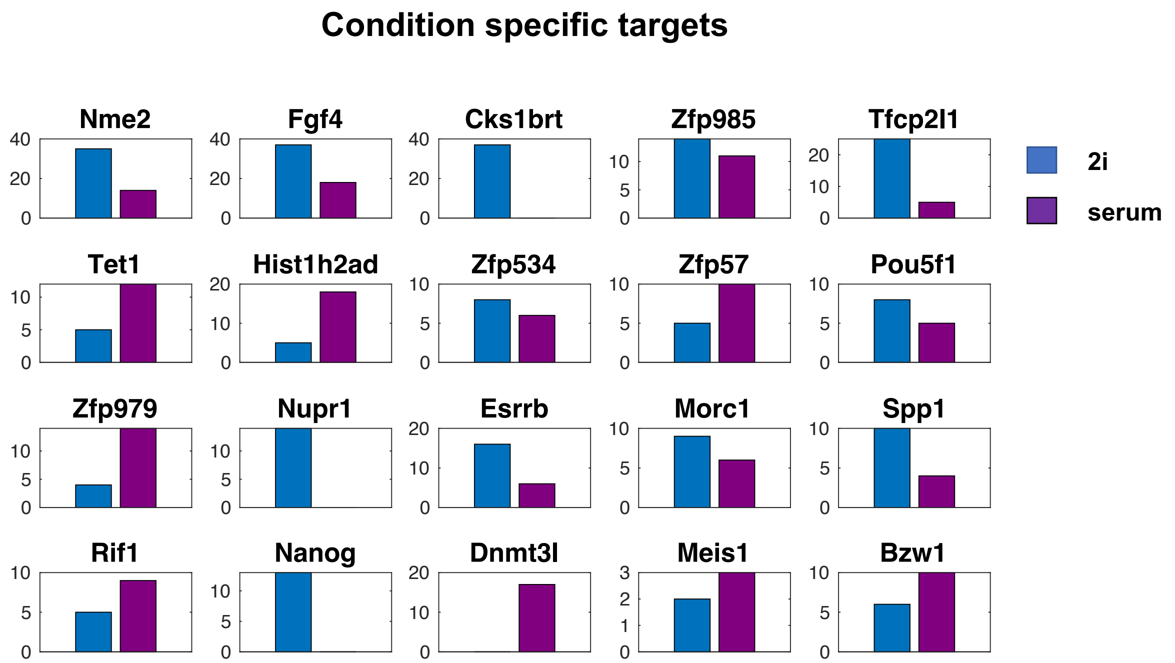


**Figure S12**. **Condition specific targets of the top 20 TFs in the mESC GRN inferred by TENET.** Nme2 and other pluripotency factors such as Fgf4, Pou5f1, and Nanog have more 2i-specific targets whereas DNA methylation factors such as Tet1 and Dnmt3l have more serum-specific targets.

network of early blood development from single-cell gene expression measurements. *Nat. Biotechnol.*, **33**, 269–276.
